# Development of a Comprehensive Antibody Staining Database using a Standardized Analytics Pipeline

**DOI:** 10.1101/563742

**Authors:** El-ad David Amir, Brian Lee, Paul Badoual, Martin Gordon, Xinzheng V. Guo, Miriam Merad, Adeeb H. Rahman

## Abstract

Large-scale immune monitoring experiments (such as clinical trials) are a promising direction for biomarker discovery and responder stratification in immunotherapy. Mass cytometry is one of the tools in the immune monitoring arsenal. We propose a standardized workflow for the acquisition and analysis of large-scale mass cytometry experiments. The workflow includes two-tiered barcoding, a broad lyophilized panel, and the incorporation of a fully automated, cloud-based analysis platform. We applied the workflow to a large antibody staining screen using the LEGENDScreen kit, resulting in single-cell data for 350 antibodies over 71 profiling subsets. The screen recapitulates many known trends in the immune system and reveals potential markers for delineating MAIT cells. Additionally, we examine the effect of fixation on staining intensity and identify several markers where fixation leads to either gain or loss of signal. The standardized workflow can be seamlessly integrated into existing trials. Finally, the antibody staining data set is available as an online resource for researchers who are designing mass cytometry experiments in suspension and tissue.

## 1 Introduction

Immune monitoring (IM) is a systems biology approach for the quantitative evaluation of the state of the immune system (Brodin and Davis, 2017, Kaczorowski et al., 2017). Changes in hematopoietic cell subset composition and in the cytokines and other proteins these cells produce can indicate the nature and severity of the stress the body is confronting. These immune correlates establish measurable proxies to the hidden details of disease or the effects of treatment, and are promising to become a central component of clinical research (Kohrt et al., 2016). Mass cytometry, which can measure over forty parameters per single cell (Bandura et al., 2009, Bendall et al., 2011), has potential applications for IM in a wide variety of contexts, including cancer (Lavin et al., 2017), allergy (Tordesillas et al., 2016, Rust and Wambre, 2017), infectious diseases (Hur et al., 2014, Cliff et al., 2015, Hamlin et al., 2017, Michlmayr et al, 2018), trauma (Seshadri et al., 2017), organ transplantation (Mario et al., 2014, Shipkova et al., 2016) and neonatal development (Olin et al., 2018). Furthermore, there is growing interest in incorporating mass cytometry into large studies such as clinical trials through the Cancer Immune Monitoring and Analysis Centers (CIMAC) and Partnership for Accelerating Cancer Therapies (PACT) initiatives^1^.

Any large-scale study will introduce challenges such as sample quality control, batch effects, and inter-operator variability. There are a plethora of methods to address potential data quality issues in mass cytometry. These include the incorporation of normalization beads into the sample (Finck et al., 2013), reduction of technical variability and doublets through multi-sample barcoding (Zunder et al., 2015, Fread et al., 2017), measurement of batch effects using spiked-in references (Kleinsteuber et al., 2016), compensation of signal spillover across different masses (Chevrier et al., 2018), and others. However, despite the well-developed ecosystem, there is no clear standard on how to run a large-scale mass cytometry study, and researchers are often forced to reinvent the wheel by designing experiments de novo with no clear guidance on best practices.

The situation is even more problematic in the computational biology arena. Numerous mass cytometry analysis methods have been published. These can be broadly classified into one of two categories. Clustering algorithms, such as SPADE (Qiu et al., 2011), PhenoGraph (Levine et al., 2015) and FlowSOM (van Gassen et al., 2015), group cells together based on marker expression patterns. Dimensionality reduction algorithms, such as t-SNE (van der Maaten and Hinton, 2008, Amir et al., 2013), embed the single cell data in a two-dimensional map that can be more easily visualized. These approaches require the operator to review their output and label cells based on his or her judgement. Despite the existence of automatic methods (Aghaeepour N et al., 2016), attempts to provide streamlined analysis workflows (Nowicka et al., 2017) and online tools such as Cytobank (PMID: 24590675), identifying appropriate analysis methods in large scale IM studies remains a challenge, and many users resort to manual gating (van Lochem et al., 2004), which is time consuming, error prone, susceptible to operator bias, and not easily scalable.

Finally, the insights gained from mass cytometry ultimately depend on the antibodies used in a given staining panel, and as with any other antibody-guided assay, antibody selection is a central component of mass cytometry experiment design. While there is some consensus on appropriate markers to identify major circulating immune subsets (Maecker et al., 2012), much of the potential of mass cytometry is in its ability to characterize the roles of less-studied markers (Horowitz et al., 2013, Bengsch et al., 2018, Good et al., 2018) and, by extension, in identifying relevant biomarkers for immunotherapy. However, there have been no systematic studies of the expression of a broad set of markers across a broad set of cell subsets to help guide antibody selection in IM studies. This problem is further exacerbated for studies involving fixed samples, since fixation can alter surface epitopes and unpredictably change antibody expression patterns (Dzangué-Tchoupou et al., 2018). A comprehensive catalog of antibody staining expression patterns across immune cells would represent a valuable resource to establish a starting point for marker selection and panel design.

In order to address the above, we developed a streamlined mass cytometry pipeline that combines a lyophilized antibody panel, two-tier barcoding, efficient batched sample acquisition and a novel cloud-based analytics service. We applied this efficient sample and data processing pipeline to screen the expression of 326 antibodies across all major peripheral blood mononuclear cell (PBMC) subsets from multiple donors on both fresh and fixed cells. This represents one of the largest mass cytometry data sets to date, with approximately 63 million events acquired over a month of operation. The workflow incorporates multiple mechanisms that address and monitor intra- and inter-sample variability, quality control, standardization and automation. The result is a comprehensive antibody staining data set, which screens marker expression in every major immune subset on a single-cell level. These antibody expression data have been made available as an interactive companion website at https://www.astrolabediagnostics.com/antibody_staining_data_set. This represents a powerful resource that allows researchers to quickly identify potential markers for inclusion in novel mass cytometry studies. Finally, the overall workflow represents a systematic framework that can readily by applied for performing IM in large experiments such as clinical trials.

## 2 Materials and Methods

### 2.1 Samples and processing

Peripheral blood mononuclear cells (PBMCs) for the primary LEGENDScreen experiment were isolated by Ficoll gradient centrifugation from leukapheresis products derived from 3 independent de-identified donors (New York Blood Center). Additional validation experiments used blood collected from consented healthy donors under an existing IRB protocol at the HIMC. For the primary screen experiment, approximately 120 million cells from each donor were incubated for 20 minutes at 37°C in RPMI media containing 10% FBS, 1μM Rh103 to label dead cells and 50μM IdU to label actively cycling cells. The samples were then washed, Fc-blocked (FcX, Biolegend) and stained for 30 minutes on ice with a lyophilized core antibody cocktail comprised of markers to allow identification of all major immune subsets (Supplementary Table 1). All the antibodies in the core panel were conjugated in-house using X8 MaxPar conjugation kits (Fluidigm), and the titrated panel was lyophilized and dispensed as single test aliquots (Biolyph). The reconstituted panel was filtered through a 0.1 micron Amicon filter prior to use.

After staining, the samples were then divided into two aliquots, one of which was fixed with freshly diluted 1.6% formaldehyde in PBS for 20 mins, while the other was left untreated. Each of the 6 samples was then barcoded using a combinatorial CD45-based barcoding scheme (Figure 1), allowing the 6 treatments to be combined as a single sample. This pooled sample of ∼300 million cells was then evenly distributed across each of the 372 wells of a LEGENDScreen kit (BioLegend) containing reconstituted PE antibodies (Supplementary Table 2), and incubated for 30 minutes on ice. Cells from each well were then washed and fixed with 1.6% formaldehyde in PBS for 20 mins. To reduce the overall number of samples to facilitate subsequent processing and data acquisition, the samples were washed with barcode permeabilization buffer (Fluidigm), and sets of 10 wells were barcoded and pooled using a combinatorial palladium-based barcoding strategy (Figure 1) (Behbehani et al., 2014, Zunder et al., 2015). The pooled samples were then washed and stained with saturating concentrations of 165Ho-conjugated anti-PE antibodies. The samples were then washed and incubated in freshly diluted 2.4% formaldehyde containing 0.02% saponin, 125nM Ir intercalator (Fluidigim) and 300nM OsO4 (ACROS Organics) for 30 minutes. The samples were then washed, frozen in FBS containing 10% DMSO and stored at −80oC until acquisition.

**Figure 1.**
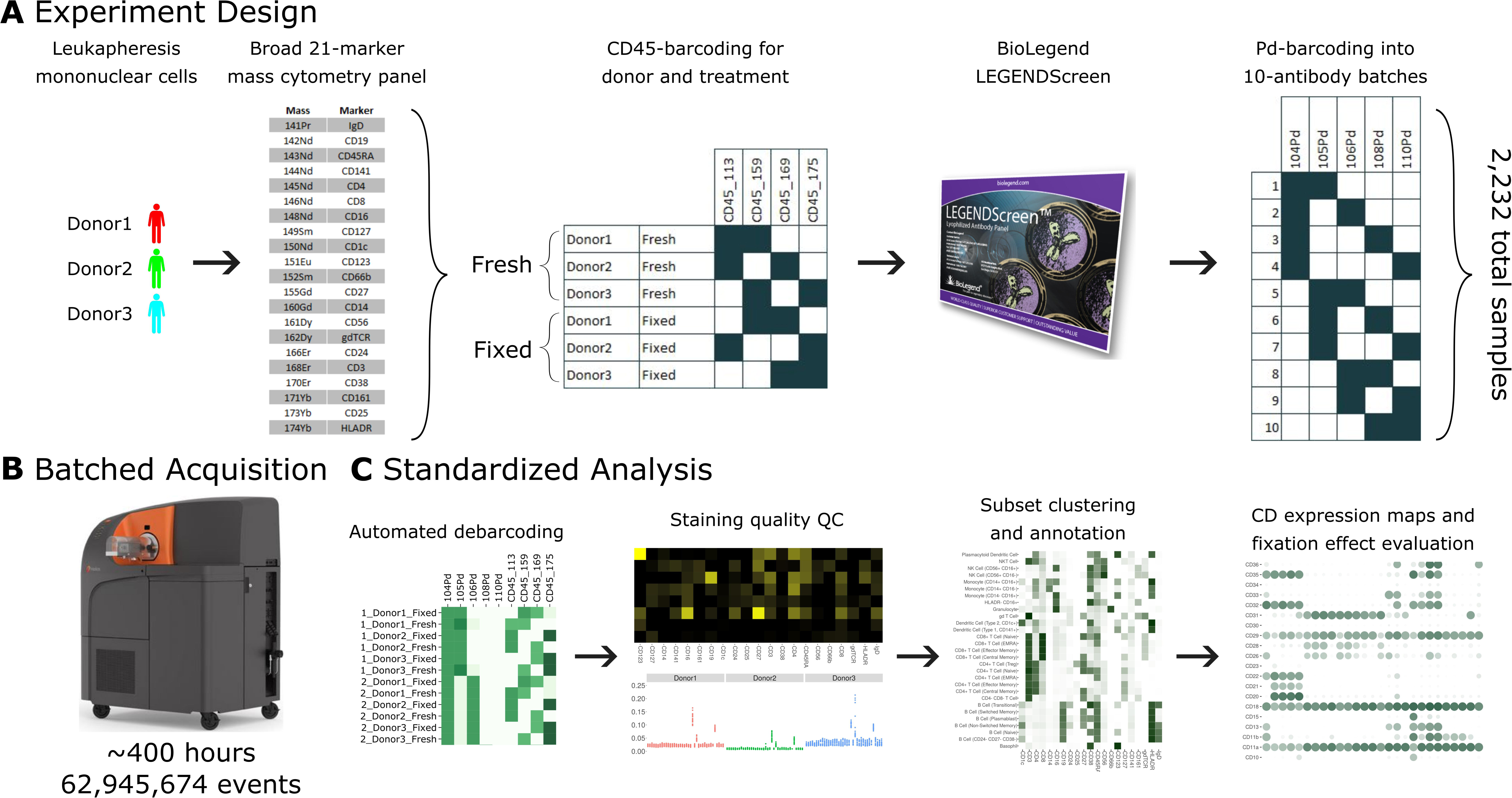
A standardized workflow for mass cytometry experiments and its implementation in generating a comprehensive antibody staining reference. A: Blood was acquired from three healthy donors and stained with a lyophilized panel of 21 metal conjugated antibodies to allow identification of major immune cell types. The samples were split into two treatments, fresh and formaldehyde-fixed. Each donor and treatment pair was barcoded using a combination of two out of four CD45 channels. Samples were divided between the four, 96-well plates of the LEGENDScreen antibody panel. Finally, the antibodies were organized into batches of ten samples each, which were in turn barcoded using a combination of two out of five palladium channels. B: The 38 batches were acquired using a Helios instrument over a period of five weeks, leading to approximately 63 million events. C: Samples were automatically debarcoded and tested for quality control using the Average Overlap Frequency (AOF), and immune populations were clustered, annotated, analyzed and visualized using the Astrolabe Cytometry Platform.

### 2.2 Data acquisition and initial data processing

Samples were thawed immediately prior to acquisition, washed once in PBS, once in CAS buffer (Fluidigm) and then resuspended in CAS buffer containing a 1/20 dilution of EQ normalization beads (Fluidigm). Following routine instrument tuning and optimization, the samples were run at an acquisition rate of <300 events per second on a Helios mass cytometer (Fluidigm) modified with a wide-bore injector (Fluidigm). Upon completion of the acquisition, FCS files associated with each barcoded batch of wells were concatenated and normalized using the bead-based normalization algorithm in the Fluidigm software resulting in 38 FCS files.

### 2.3 Mass cytometry data analysis

FCS files were uploaded to the Astrolabe Cytometry Platform (Astrolabe Diagnostics, Inc.) where transformation, debarcoding, cleaning, labeling, and unsupervised clustering was done. Data was transformed using arcsinh with a cofactor of 5 and the marker intensities presented in the paper are all after transformation. Batches were debarcoded using the Ek’Balam algorithm (see below), resulting in 2,232 individual samples corresponding to one (donor, treatment, antibody) combination. Data from twelve antibody wells were excluded due insufficient cell recovery or ambiguous barcoding resulting from a known pipetting errors during sample preparation, resulting in 2,160 samples. For batches 23, 25, and 34, between 50% and 75% of events were removed due to loss of stability, as described in the main text.

The individual samples were then labeled using the Ek’balam algorithm (Supplementary Table 3). Each cell subset was clustered using the profiling step in Astrolabe (see below). For the purpose of the Ek’balam algorithm, gdTCR intensities were compensated by 1.9% of CD8 intensity due to known signal spillover due to oxide formation from the 146Nd-CD8 channel being detected in the 162Dy gdTCR channel. Platform output was downloaded in the form of R Programming Language RDS files (R Core Team, 2013) for manual follow-up analysis. Figures were generated using ggplot (Wickham, 2016). To evaluate the quality of the the debarcoding, clustering and annotation in Astrolabe and to perform independent analyses, a subsets of samples were processed in parallel using a Matlab based debarcoding algorithm (Fread et al., 2017) and uploaded to Cytobank for manual gating of major immune subsets.

### 2.4 The Ek’Balam algorithm

Ek’Balam is a hierarchy-based algorithm for labeling cell subsets which combines the strength of a knowledge-based gating strategy with unbiased clustering. It receives a user-defined subset hierarchy which details gating rules such as “Cells which are CD3+ are T Cells”. Subsets can branch through additional rules, for example, “T Cells which are CD4+ are CD4+ T Cells”. The hierarchy is organized into levels which correspond to parallel steps when gating. For example, the first level could include “CD3+ are T Cells”, “CD19+ are B Cells”, and “CD33+ are Myeloids”. Ek’Balam then iterates over the levels. At each iteration, the data is clustered with FlowSOM (van Gassen et al., 2015), using only the markers that appear in the rules of that level. Each cluster is then labeled according to the rules of that level. Labeling is done by optimizing the Matthews Correlation Coefficient (MCC) over the clusters and marker intensity values with a greedy algorithm. The process continues until all cells are assigned to a label which has no rules branching out of it. A formal definition of the algorithm is provided in the supplement.

### 2.5 Cell subset profiling

Profiling refers to a variation of unsupervised clustering using the FlowSOM algorithm. The variant differs from classic FlowSOM in two significant aspects. One, each cell subset is clustered separately. This guarantees that the output will not include biologically irrelevant clusters that combine multiple cell subsets. Two, the clusters are labeled according to the markers that differentiate between them the most, according to the MCC. The labeling makes the output more accessible to the researcher by providing an initial intuition about the differences between the clusters. A formal definition of the profiling algorithm is provided in the supplement.

### 2.6 Relevance metrics

The following metrics were employed when comparing the computational debarcoding and labeling results to manual methods. Metrics were calculated for each class separately, where class is either a barcode (for debarcoding) or a cell subset (for labeling). The class was set as the target and all other classes as not-target. In all cases, the manual method is assumed to be the correct solution.

TP, FP, TN, and FN are true positive, false positive, true negative, and false negative, respectively.

Precision is the frequency of correctly classified target events out of all events classified as target, or TP / (TP + FP).

Recall is the frequency of correctly classified target events out of all target events, or TP / (TP + FN).

The F1 score is the harmonic mean of precision and recall, or

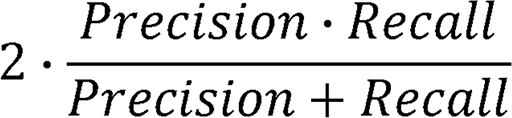

The Matthews Correlation Coefficient (MCC) is the correlation coefficient between the computational and manual classification, or

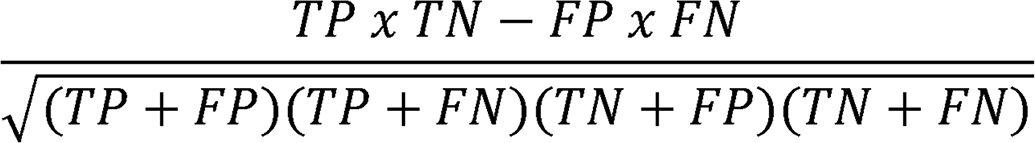

### 2.7 Average overlap frequency (AOF)

The average overlap frequency is a metric of staining and clustering quality of a given marker (Amir et al., 2018). It assumes that the marker has two modalities, denoted negative and positive. The AOF is a value between 0 and 1, where 0 is complete separation between the modalities and 1 is complete overlap, and is defined as:

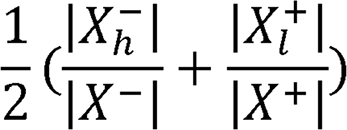

where *X*^-^ is the values of all events in the negative modality, *X*^+^ is the values of all events in the positive modality, 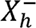 is the negative values that are greater than the 5th percentile of a normal distribution with a mean and standard deviation of, and 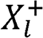 is the positive values that are lower than the 95th percentile of the normal distribution with mean and standard deviation of.

Given a set of samples, we can extend the AOF into a sample quality score by calculating the *Scaled*^*2*^*AOF* for each (marker, sample) pair:

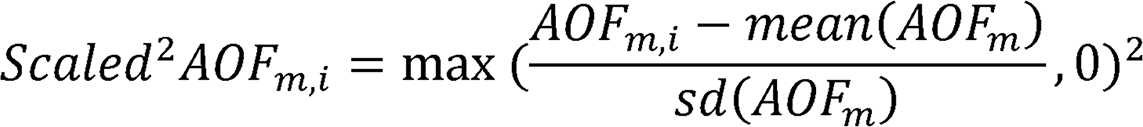

where m indexes over markers and i indexes over samples, and then calculating the *Quality*^*2*^*AOF* for each sample:

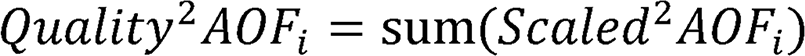

### 2.8 Percent positive events

For each (profiling subset, antibody), the percent of positive events is the percent of events whose intensity is greater than the 99th percentile of all events in the Blank LEGENDScreen well (well A1 in plate 1, see Supplementary Figure 3). This well does not include any PE-conjugated antibodies, so the intensity distribution there is a background for anti-PE measurement using the Helios. In order to assess the potential effect of the isotype control on the baseline, we calculated an alternative percent positive based on the 99th percentile of the respective isotype for each antibody. The correlation between the Blank-based and the isotype-matched percent positive values was 0.94 and the median different was 1%. Due to this minor difference we decided to use the same Blank 99th percentile for all antibodies.

**Figure 3.**
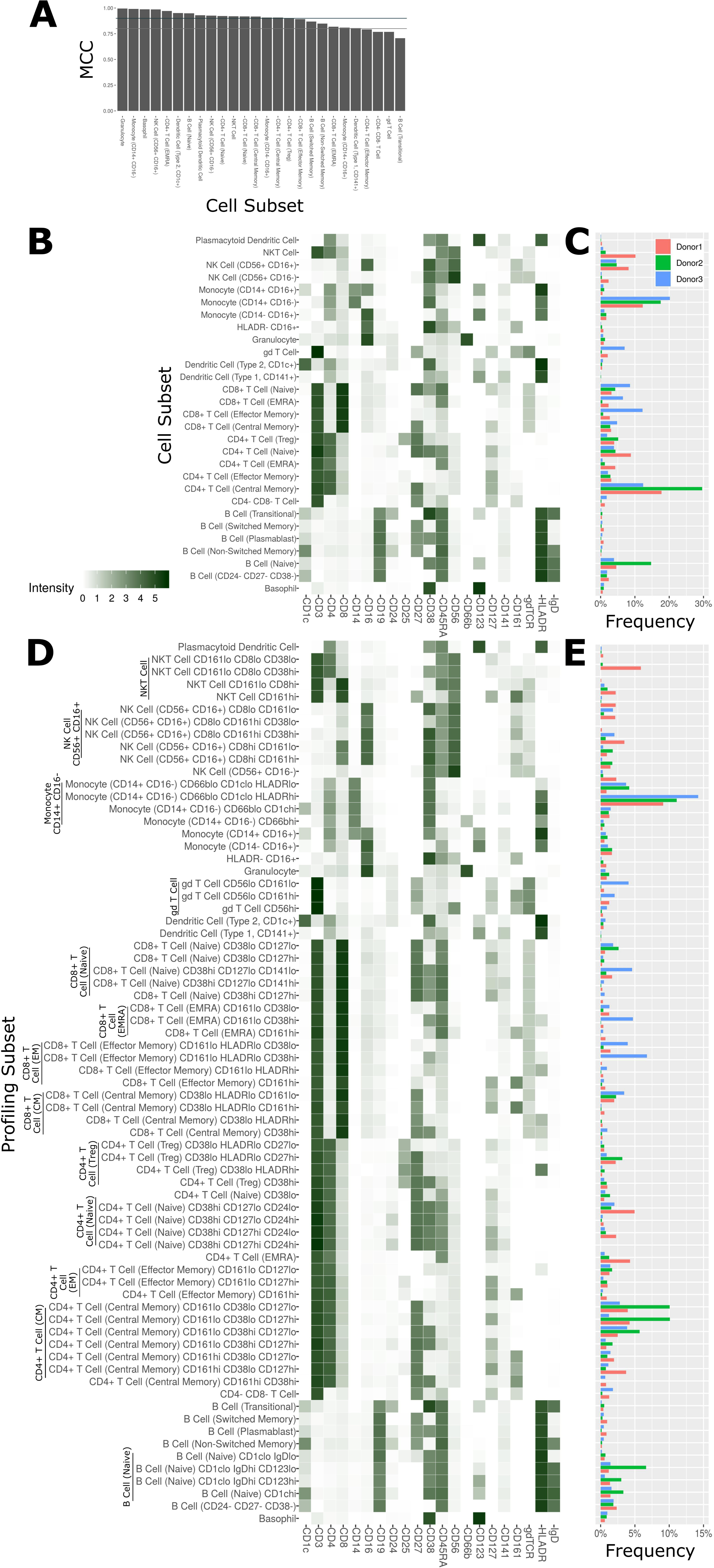
Cell subset and profiling subsets using the Astrolabe platform. Cells were clustered using the FlowSOM algorithm and labeled according to canonical gating hierarchy. A: Bar plot comparing Astrolabe labeling to manual gating across all cell subsets using the Matthews Correlation Coefficient (MCC). Jade line corresponds to MCC of 0.9, grey line to MCC of 0.8. Almost all cell subsets have a high MCC, denoting agreement between the automated and manual methods. B: Cell subset heat map for one example sample. Rows are cell subsets, columns are channels. Tile intensity is median of channel in subset. C: Bar plot of mean cell subset frequencies in each donor. D: Profiling subset heat map for the same example sample. E: Bar plot of mean profiling subset frequencies in each donor.

## 3 Results

### 3.1 Design of an Integrated Pipeline for the Acquisition and Analysis of Large Immune Monitoring Experiments

Conducting a large-scale immune monitoring experiment over a long period of time using mass cytometry raises several challenges. One, it is imperative to monitor instrument performance and evaluate sample data quality to identify transient fluctuations in instrument performance resulting in features such as diminished staining for one or more markers, higher than usual debris or doublet count. Two, batch effects due to experimental or instrument variation can be a significant concern. While researchers should always be aware of how technical sources could lead to variation, this is especially pertinent when data is gathered and acquired over weeks or months. Experiment design should therefore include mechanisms that detect both types of failures and alert the researcher appropriately. Finally, the role of human operators should be minimized in order to reduce human-introduced variability. Decision making should follow a clear protocol or be entrusted to computational methods.

The antibody expression data set described in this study integrates multiple techniques to minimize experimental and technical reproducibility and streamline data acquisition and analysis (Figure 1). Peripheral blood mononuclear cell (PBMC) samples from three healthy donors (Figure 1A) were stained with a 21-marker antibody panel comprised of markers to unambiguously identify all the major immune compartments: B Cells, myeloid cells, NK Cells, and T Cells, together with further granularity for subsets within these compartments (such as CD16+/- monocytes or naive versus transitional B Cells). This core antibody panel was lyophilized as a single cocktail and the same batch was used throughout sample acquisition to minimize experimental variability due to reagent variability or pipetting. The panel only utilizes a subset of the channels available in mass cytometry, allowing researchers to incorporate an additional 10-15 markers to address experiment-specific questions.

Following initial core antibody panel staining, the samples were split into two groups to evaluate the impact of fixation on each of the antibody epitopes subsequently evaluated in this screen. This design also typifies a common experimental design where a treatment (fixation) is compared to control (fresh samples). The six patient x treatment combinations were barcoded and pooled using a live cell-compatible doublet-free barcoding strategy leveraging CD45 antibodies conjugated to 4 distinct isotopes. This barcode approach streamlines sample processing and minimizes potential variability due to acquiring different patients or treatments at different times. The isotopes used for barcoding were specifically chosen to ensure that potential spillover due to isotopic impurities or oxide formation from these barcoding channels would not influence any of the other antibody channels being measured in this experiment. Next, the samples were evenly distributed across each of the 372 wells of a LEGENDScreen kit, each of which includes a PE-conjugated antibody against a distinct epitope. Following this with a metal-conjugated anti-PE antibody enabled the measurement of a comprehensive set of surface markers across all the cell subsets identified by the broad lyophilized panel. Finally, to streamline data acquisition, sets of 10 wells were further barcoded and combined using a combinatorial strategy leveraging five palladium channels.

The resulting 38 batched samples were then acquired using a Helios mass cytometer (Figure 1B). Acquisition required around 400 hours of instrument time over five weeks of operation and resulted in a total of 63 million events. Analyzing such a large amount of data manually would have been time-consuming and risked operator-introduced variability. To avoid these two issues, we employed the standardized Astrolabe Cytometry Platform to debarcode and clean the data, label cell subsets, and conduct unsupervised clustering (Figure 3C). The Astrolabe analysis took 24 hours, and the platform’s “Analysis” export was employed in all follow-up analyses.

### 3.2 Debarcoded Sample Data is Robust and Consistent Across the Screen Samples

The antibody staining data set involves a high number of samples, complex experiment design, a long acquisition period, and advanced computational analysis, any of which could potentially introduce variability or other artifacts. Several tests inspect the various stages of the experiment (Figure 2). First and foremost, accurate debarcoding is critical for all follow-up analyses. This step is especially challenging due to the two-tiered barcoding scheme employed: CD45-based barcoding of patient x treatment and palladium-based barcoding of each batch of 10 LEGENDScreen antibodies. Astrolabe correctly identifies all 60 codes and their channel profile are distinct and follow the expected design (Figure 2A). In order to validate the computational debarcoding approach, the results were compared to manually-debarcoded data for one of the batches. The two methods showed high concordance according to four different statistical metrics (Figure 2B), supporting the use of the more efficient computational approach to debarcode all 2,232 samples.

**Figure 2.**
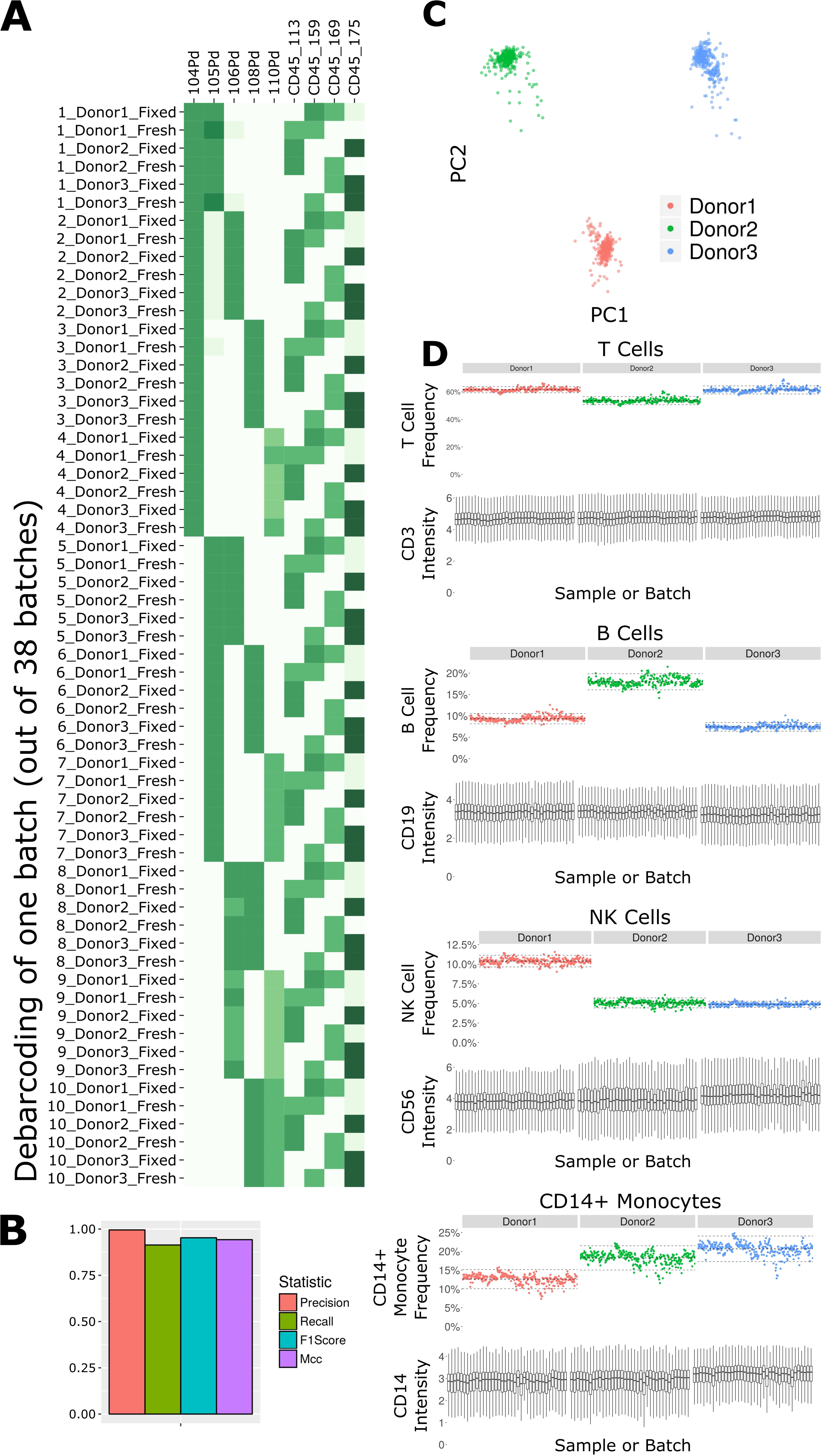
Subset frequency and marker intensity are consistent across the LEGENDScreen experiment acquisition. A: Debarcoding heat map for one batch out of the 38. Rows are debarcoded events, columns are barcoding channels. Tile intensity is median of channel in events. Astrolabe correctly debarcoded all of the codes in this batch. B: Bar plot comparing Astrolabe debarcoding to manual debarcoding using four accuracy scores: Precision (0.996), Recall (0.914), F1 Score (0.953), and Matthews Correlation Coefficient (MCC, 0.943). C: Scatter plot of Principal Component Analysis (PCA) over the cell subset frequency vectors for all samples. Axes are first and second components, dots are samples color-coded by donor. The three donors are distinct from each other and internally uniform across the entire acquisition. D: For each pair, the top plot is a scatter plot of cell subset frequency across all samples, ordered by batch. The bottom plot is a box plot of the canonical marker intensity across all batches. From top to bottom: T Cells and CD3 intensity, B Cells and CD19 intensity, NK Cells and CD56 intensity, and CD14+ Monocytes and CD14 intensity. In all cases, both frequencies and intensities are robust across the acquisition period.

The starting point for the data set was blood from three healthy donors. After the fixed versus fresh treatment and the introduction of the kit’s antibodies, each of these individuals leads to several hundred different samples. However, the individual donor immune profile across each set of samples are expected to be identical and therefore the acquired data should be highly comparable. This is reflected in the principal component analysis (PCA) map over the sample cell subset frequencies (Figure 2C). The samples are distributed across three well-separated islands. Each island corresponds to one individual, signifying that the immune profile is consistent throughout acquisition.

We further applied Average Overlap Frequency (AOF) as a metric to evaluate individual marker staining quality across all sample batches (Amir et al., 2018). This QC step identified issues with staining of multiple markers in three of the batches (Supplementary Figure 1A). Further inspection of the score highlighted several problematic markers (Supplementary Figure 1B). Evaluation of the single-cell data for one of these markers, CD27, revealed a time-dependent increase in background staining resulting in reduced marker resolution over time, which we attribute to a Helios instrument malfunction during acquisition (Supplementary Figure 1C). However, restricting analysis to only the events in the first quarter of acquisition window for these batches resulted in AOF values within the range of other batches, allowing recovery of valid antibody screening data despite the technical issues (Supplementary Figure 1D). The rapid identification, isolation, and solution of these technical artifacts was facilitated by a standardized quality control approach using the well-defined AOF metric.

Except for the batch effects identified by the AOF QC, the data set was consistent across cell subsets and marker intensities (Figure 2D). For four major cell subsets (from top to bottom: T Cells, B Cells, NK Cells, and CD14+ Monocytes), we examined the frequency in each sample (top panel of each, ordered by batch). Subset frequency has very small variation across all the samples of a given donor. Additionally, the distribution of the canonical marker of each subset (CD3, CD19, CD56, and CD14, respectively) is also consistent across the samples (bottom panel of each, one box for each batch).

The combination of the above quality control measures highlights the overall robustness of the antibody staining data set. The overall staining data were cohesive for each donor, and for each cell subset across donors, and specific acquisition issues were identified and addressed using automated QC metrics.

### 3.3 The Astrolabe Platform Correctly Labels Cell Subsets and Provides Meaningful Unsupervised Clustering

The Astrolabe platform automatically labeled canonical immune cell subsets (Figure 3). As with debarcoding, it is imperative to verify that automated cell annotation methods correspond to historical definitions by calculating the overlap with manual gating. The Matthews Correlation Coefficient (MCC) between the two methods was greater than 0.8 for almost all of the cell subsets (Figure 3A). Biaxial plots of canonical markers further reinforced the overlap (Supplemental Figure 2A-C). Four of the subsets had a score lower than 0.8, which indicated some discrepancy between computational labeling and manual gating. In all four cases, the disagreement was due to subjective thresholding of a specific marker (Supplementary Figure 2D): these are cases where the exact marker intensity threshold for a given subset is ambiguous, such as where to draw the line on CD24 to distinguish Naive and Transitional B Cells. Importantly, the automated approach allowed consistent thresholding across all samples in these ambiguous cases, avoiding potential human subjectivity and variability in assigning gates across samples.

The marker intensity profiles for each of the subsets labeled by the platform largely follow the consensus HIPC definitions (Figure 3B, Maecker et al., 2012). Astrolabe consistently identified 11 T Cell subsets (including CD4+ and CD8+ T cells, and Naive, EMRA, EM and CM subsets within each), 6 B Cell subsets, several myeloid subsets, NK Cell subsets, granulocytes, and NKT Cells. Examining cell subset frequencies across the three donors highlighted clear variability in their respective immune profiles (Figure 3C), which further reinforces the previous PCA results.

The discovery of novel cell subsets defined by previously unappreciated marker expression patterns is one of the most exciting promises of high-complexity cytometry such as mass cytometry. While cell subset labeling follows established trends, unsupervised clustering has the potential to unearth previously unknown signals. Astrolabe includes a profiling step, where each defined cell subset is clustered separately (Figure 3D). The number of clusters is decided via a heuristic which depends on the number of cells in each subset and on marker heterogeneity. In the antibody staining data set, the platform returns 71 profiling subsets, which are then labeled according to the marker or markers that provide the greatest separation between them. Notably, several CD8+ T Cell subsets are broken down based on CD161, suggesting MAIT-like T Cells (Ussher et al., 2014). Naive B Cells are differentiated based on IgD, while NK Cells are broken up according to CD8. Similar to the canonical cell subsets, profiling subset frequencies vary between the three donors (Figure 3E), hinting at a wider heterogeneity within the population.

### 3.4 The Antibody Staining Data Set Defines Expression Patterns of Hundreds of Surface Markers across 71 Cell Subsets

With 350 measured antibodies over 71 profiling subsets, the antibody staining data set is a rich source of information about expected expression patterns in a healthy immune system. In order to provide an initial view into the full expression dataset, we calculated two metrics for each profiling subset and antibody combination (Figure 4A, Supplementary Figure 3). The first metric is the median marker intensity, which is most useful in defining expression of markers that show a unimodal distribution within a given subset. To better reflect bimodal expression patterns, or those in which only a subset of cells are positive for a given marker, we used a blank well that lacked any PE-primary antibody to establish a baseline for the second metric, percent positive cells. We set an arbitrary cutoff at the 99th percentile of the blank well and defined any cell above this value as positive for the marker. The resulting heat map provides two separate summary statistics of marker expression over all profiling subsets.

**Figure 4.**
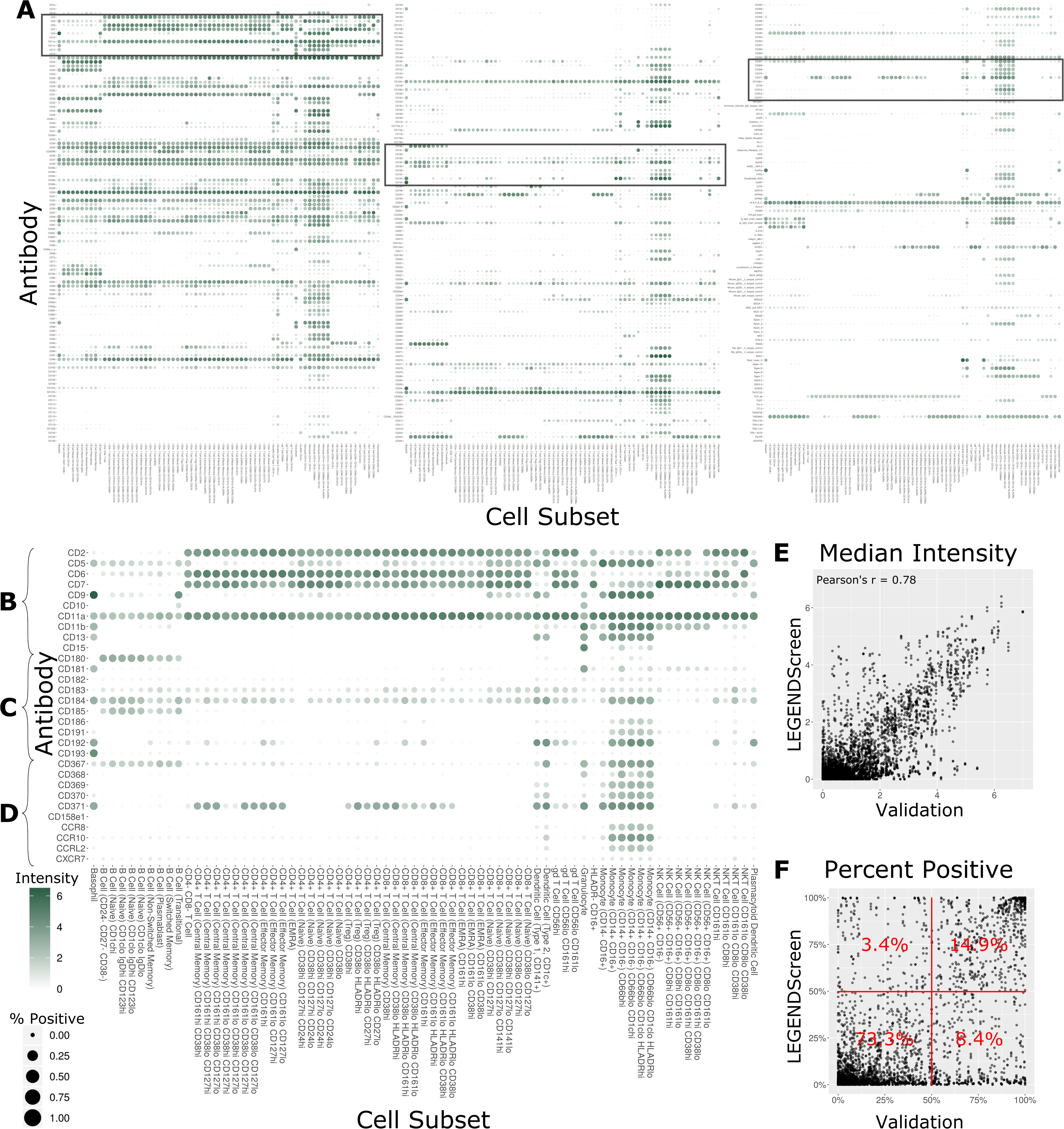
Summary of LEGENDScreen antibody intensities across all profiling subsets. A: Heat map of all antibodies across all profiling subsets. Rows are antibodies, columns are profiling subsets. Tile size is percent positive and intensity is median intensity. The three highlighted sections correspond to panels B, C, and D. B, C, and D: Larger view of these sections of the main heat map. Sections were chosen to broadly correspond to the top, middle, and bottom of the heat map. E: Scatter plot comparing median antibody intensity between the LEGENDScreen experiment and an independent validation experiment using a fourth donor sample that was processed and acquired separately. X-axis is median intensity in validation, Y-axis is median intensity in LEGENDScreen. Each dot corresponds to one cell subset and antibody combination (Pearson’s rho is 0.78). F: Scatter plot comparing percent positive between experiments. Percentages are frequency of markers in each quadrant (McNemar’s Chi-squared test p-value is lower than 2.2-16).

Focusing on any specific section of the heat map reveals a plethora of relevant patterns. The top of the map is populated with well-established markers (Figure 4B) such as CD7, which is present on all T Cell and NK Cell profiles, and CD11b, which is most highly expressed by monocytes. This section also highlights a limitation of the data set with CD5: while this is generally considered a pan-T cell marker the screen only showed expression on Naive CD4+ T Cells, and not any other CD4+ T Cells. This idiosyncratic staining pattern could be due to many potential reasons, such as limitations of the LEGENDScreen kit, antibody clone used, or specifics of the Helios protocol that we employed. This serves as an important reminder to researchers who are looking to utilize this resource: as with any other biological screen, specific signals should be further validated before being relied upon.

Lower sections of the heat map allow investigation of many surface markers that appear less frequently in the scientific literature (Figure 4C). Notably, the screen reproduces the expression of CD180 in B Cells (Miura et al., 1998) and the expression of CD193 in basophils (Florian et al., 2006), while revealing new potential patterns such as the expression of CD181 by granulocytes and basophils. Additionally, many markers are expressed by myeloid cell subsets to some degree. It remains to be seen whether this is an artifact of the experimental technique employed here, or whether there is a high degree of myeloid cell heterogeneity that still remains to be defined. This trend continues throughout the heat map (Figure 4D), as are some more elusive signals, such as CD371, which has a checkered expression pattern across diverse and seemingly unrelated profiling subsets.

In order to provide some outside validation for the dataset, we conducted a second independent LEGENDScreen experiment using PBMCs from a fourth donor and compared marker medians (Figure 4E) and percent positive (Figure 4F) between the two experiments (Supplementary Table 4). The metrics are correlated between the experiments (Pearson’s rho of 0.78 and 0.73, respectively). For percent positive, 88.2% of markers are in the bottom left or top right quadrant of the plot, showing high agreement between the original experiment and validation experiment (McNemar’s Chi-squared test p-value is lower than 2.2-16). Together, these tests show that the trends seen in this data set are generalizable. With that said, unlike the main data set, CD5 is uniformly expressed across all T Cell subsets in the validation data (Supplemental Figure 4), further reinforcing the importance of validation of screens. Examining the set of markers that are distant from the diagonal does not reveal any clear trends and it is possible that they are a result of donor-specific differences, technical variation between the experiments, or random noise.

### 3.5 Several Markers are Differentially Expressed Between CD161+ and CD161- CD8+ T Cells

This comprehensive antibody resource offers opportunities to identify markers to further interrogate or stratify specific immune cell subsets. As a proof of principle of this approach, we leveraged the inclusion of CD161 in the core antibody staining panel, a marker that is highly expressed on mucosal associated invariant T (MAIT) cells (Martin et al., 2009). MAIT cells are a subset of T cells that display innate-like qualities (Napier et al., 2015), including an invariant TCRα chain (Porcelli et al., 1993) and an inherent capacity to respond to infection (Gold et al., 2013). The Astrolabe profiling identified CD161hi and CD161lo subsets for both Central Memory (CM) CD8+ T Cells and Effector Memory (EM) CD8+ T Cells (Figure 3D). These profiling subsets were further explored for differential marker expression trends (Figure 5). Comparing the percent positive metric for each antibody and looking for a consensus across all three donors identified six differentially expressed markers in CM cells (Figure 5A) and four markers in EM cells (Figure 5B).

**Figure 5.**
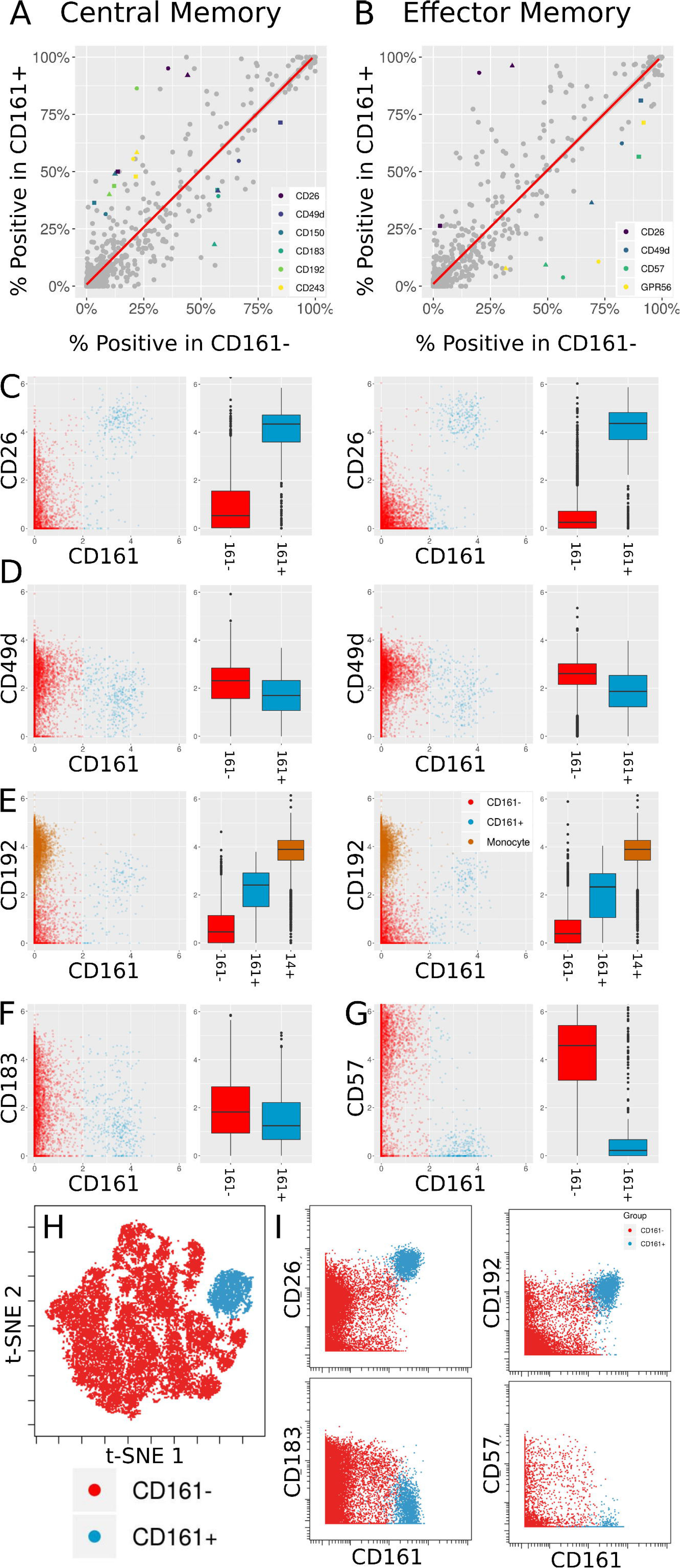
Differentially expressed markers between CD161+ and CD161- CD8+ T Cells. A, B: Scatter plots showing percent positive of each marker in different types of CD8+ T Cells. A corresponds to Central Memory (CM), B to Effector Memory (EM). X- and Y-axes are percent positive in CD161- and CD161+ cells, respectively. Each dot corresponds to one marker in one patient. The red line is linear regression. Markers where absolute standardized regression residual is greater than 2 for all three donors are colored. C: Biaxial plots and box plots of CD26 expression in CM (left) and EM (right) cells. X-axis is CD161, Y-axis is CD26, each dot is a cell. Cells and boxes are color-coded by CD161- (red) and CD161+ (blue). D: Same for CD192 expression. CD192 levels in monocytes (brown) are provided as reference. E: Same for CD183 expression in CM cells. F: Same for CD57 expression in EM cells. G: Multiplexed validation of differentially expressed proteins between CD161+ and CD161-subsets using an independent PBMC sample. tSNE map highlighting the distinct high dimensional phenotype of the CD161hi cell subset (blue). H: Biaxial plots showing validation of CD26, CD193, CD183, and CD57 expression in CD161- and CD161+ CD8+ T Cells

Two of these trends overlap between the two cell subsets: an increase in CD26 and a decrease in CD49d. CD26 has been previously associated with MAIT cells (Dusseaux et al., 2011). When examining anti-PE in the CD26 LEGENDScreen well (Figure 5C), there is a x4.5 fold increase in intensity on average between CD161- and CD161+ CM cells and a x7.2 fold increase on average between CD161- and CD161+ EM cells. For CD49d (Figure 5D), the average decrease in intensity is x1.2 and x1.5, respectively, which is to be expected given the overall low intensity for that marker.

CD192 (CCR2) was differentially expressed between CD161hi and CD161low CM cells, with a x3.6 average fold increase in intensity in the CD161hi subset (Figure 5E, left). It was only differentially expressed for two of the three donors in EM cells (figure 5E, right). CD192 is involved in recruitment of monocytes to inflammatory sites (Tsou et al., 2007), a function that could potentially be shared by MAIT cells. When examining marker intensities on a single-cell level, the CD161hi cells are situated between the CD161low cells and monocytes, and would thus be classified as CD192mid using standard gating nomenclature. In addition to these markers that were selectively upregulated on CD161hi cells, the screen highlighted reduced expression of CD183 on CD161hi CM cells (Figure 5F) and CD57 on CD161hi EM cells (Figure 5G).

One of the limitations of this screening approach is that each of the antibodies is profiled independently, which precludes co-expression analyses of markers in the screen. To validate and further explore the co-expression patterns of the markers identified in the screen, we independently stained a healthy donor PBMC sample with a panel incorporating several of the differentially expressed markers identified in the screen together with Va7.2 TCR to definitively identify MAIT cells (Supplementary Table 5). tSNE analysis on the gated CD8 T cells revealed that the CD161hi population had a distinct phenotype in high dimensional space defined by co-expression of many of the markers identified in the screen (Figure 5H and Supplementary Figure 5). The differential expression patterns of CD26, CD192, CD183, and CD57 between the CD161hi and CD161low largely mirrored those see in the initial screen, independently validating these results (Figure 5I).

### 3.6 Sample Fixation Leads to Both Loss and Gain in the Intensity of Specific Markers

Formaldehyde fixation is a useful approach to preserve cell samples but has been associated with changes in cell surface epitopes and marker expression profiles (McCarthy et al., 1994, Dzangué-Tchoupou et al., 2018, Supplementary Figure 6). However, given the prevalence and importance of fixation in cytometry experiments, there is an urgent need for a systematic study of the effect of fixation on marker intensity to better inform marker selection and panel design in studies involving fixed samples.

**Figure 6.**
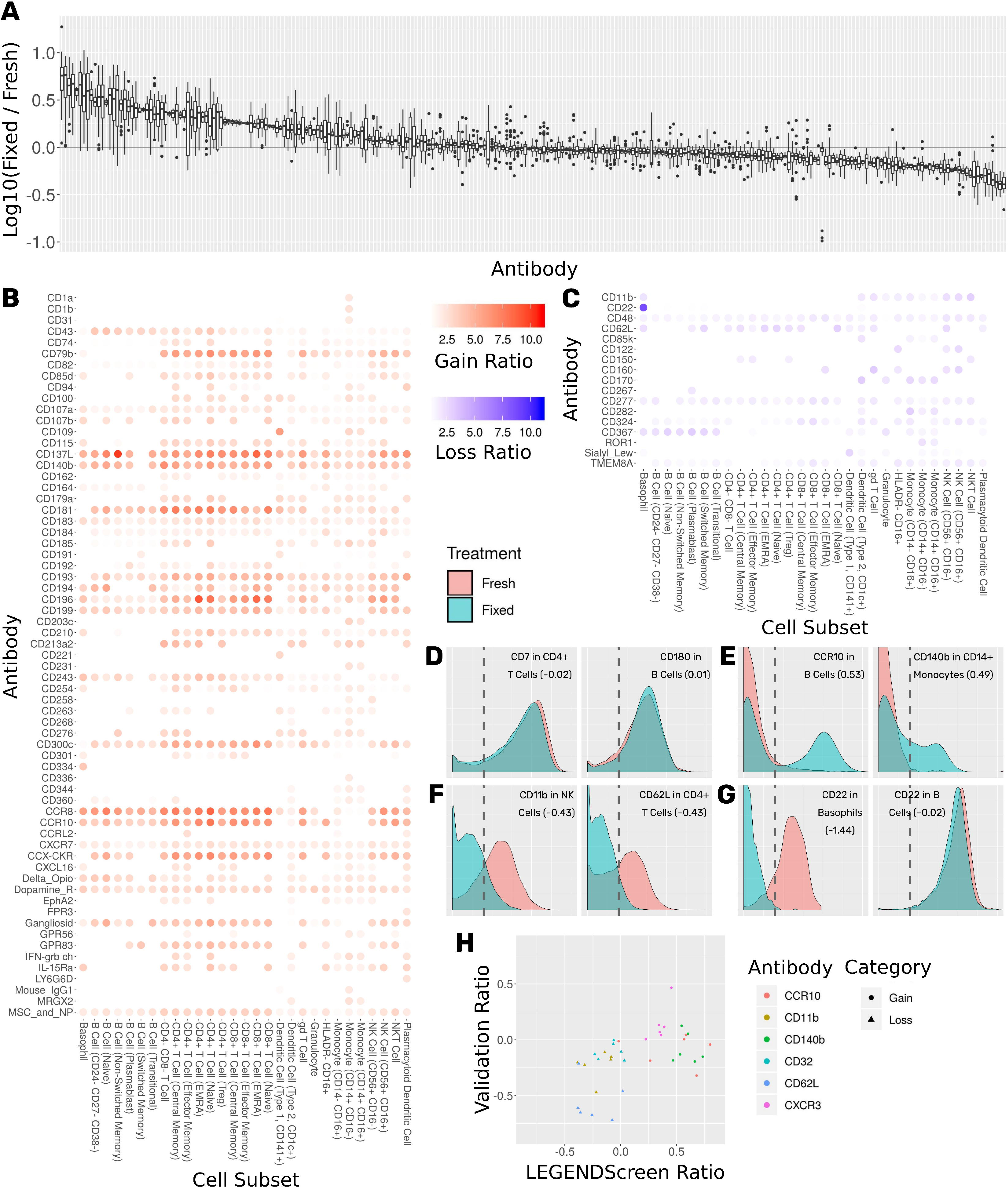
Comprehensive assessment of the effect of fixation on marker intensity. A: Box plot showing the ratio between median intensities in the fixed and fresh samples. X-axis is antibody, ordered by median ratio, Y-axis is log10 of ratio. Each box is one antibody. The grey line corresponds to zero, or equal median intensities. B: Heat map of antibodies where signal was gained from fresh to fixed (log 10 of ratio greater than 0.5). Rows are antibodies, columns are cell subsets. Tile intensity is fold change between fixed and fresh. C: Heat map of antibodies where signal was lost from fresh to fixed (log 10 ratio lower than −0.5). Tile intensity is fold change between fresh and fixed. D: Intensity distributions for two (cell subset, antibody) combinations where median was equal between fresh (in red) and fixed (in cyan). X-axis is antibody intensity, Y-axis is density. The dashed line is the 99th quantile of the blank and indicates the background. Values in parenthesis are log10(Fixed / Fresh) for this marker in this cell subset. E: Two (cell subset, antibody) combinations where median increased in fixed (gain of signal). F: Two (cell subset, antibody) combinations where median decreased in fixed (loss of signal). G: Left, CD22 antibody in Basophils, where signal was lost. Right, CD22 antibody in B Cells, where signal remained constant. Fixation effect differed between the subsets. H: Scatter plot comparing ratio of fixed and fresh between the LEGENDScreen experiment and a validation experiment where indicated antibodies were part of the mass cytometry panel (not conjugated to anti-PE). X-axis is ratio in validation, Y-axis is ratio in LEGENDScreen. Each dot is a (cell subset, antibody) combination. Color is antibody, shape is category (gain or loss).

The antibody staining data set includes two conditions for each donor and antibody samples: one stained fresh and stained following fixation with 1.6% formaldehyde. 255 of the LEGENDScreen markers have cells whose intensity is higher than the blank threshold. For each of these markers, we calculated the ratio between median expression in each of the conditions over all cell subsets (Figure 6A). We arbitrarily set a threshold of two-fold change as indicative of a significant intensity shift between the conditions. 173 (68%) of the markers were below that threshold suggesting that they are not notably affected by fixation.

Sixty-five of the markers have a two-fold or more increase in fixed samples relative to fresh (Figure 6B). In other words, these markers gained additional signal when the sample was fixed. This increase in expression can either be an artifact of fixation or true expression of an antigen that was not detected in the corresponding fresh sample. While formaldehyde fixation may be expected to partially comprise the cell membrane, the samples in this screen were not explicitly treated with any permeabilizing agents, so we do not anticipate significant exposure of intracellular antigens. Furthermore, gains in expression were largely seen across most cell subsets, suggesting that in most cases these reflect non-specific staining artifacts following fixation. At the opposite end of the spectrum, 17 markers showed a two-fold or more decrease from fresh to fixed and were thus classified as loss of signal (Figure 6C). Since only an existing signal can diminish, the lost pattern is specific to certain subsets.

Examining the ratio between the medians enables a broad survey of all antibodies over all subsets. However, it ignores the single-cell nature of the data. Closer examination of several marker intensity distributions reveals that when the ratio is around zero, the underlying distribution is usually maintained from fresh to fixed as well (Figure 6D). When marker intensity is gained, it typically only affects some of the cells within the subsets, while the low expression persists in others (Figure 6E). On the other hand, when signal is lost, it appears that fixation diminishes it completely (Figure 6F). These trends further reinforce the hypothesis that the signal gained by fixation is due to the protocol rather than the underlying biology. In almost all cases, changes in markers expression patterns showed similar trends across subsets expressing that marker. One notable expression was CD22, which was found to be expressed on both B cells and basophils in the fresh samples using the clone contained in the Legendscreen panel (S-HCL-1), consistent with previous descriptions of clone-specific CD22 expression on basophils (Han et al., 1999, Han et al., 2000). However, fixation resulted in loss of expression specifically on basophils, but not on B cells (Figure 6G), reflecting differences in the fixation sensitivity of the CD22 conformational epitopes that are differentially expressed between B cells and basophils (Toba et al., 2002).

The LEGENDScreen kit includes antibodies conjugated to PE which are then measured by mass cytometry using an anti-PE secondary. It is possible that the effects of fixation observed here are not due to effects on the underlying antibody, but rather due to a more complex interaction that potentially includes the marker antibody, PE, and anti-PE. We therefore performed a validation experiment where seven of the gain or loss markers were incorporated into the mass cytometry panel (Figure 6H). For the three loss markers, the validation results confirm the effect we saw in the data set: the same subsets express these markers, and loss their signal after fixation. On the other hand, the results for the gain markers were mixed. While one of them (CXCR3) fully reproduced the screen results, the other two only lost their signal in some of the cell subsets.

Cytometry experiment design can be a daunting task due to the high number of variables that needs to be considered. There are many factors that could influence results in unknown ways, especially when employing a method such as fixation that has the potential to perturb the chemistry and kinetics underlying the assay. This antibody staining data set represents an accessible resource to identify and anticipate such potential effects.

## 4 Discusion

We present a standardized workflow for the acquisition and analysis of large-scale immune monitoring studies using mass cytometry. The workflow incorporates several established experimental techniques in order to reduce signal variation within samples, across samples, and across operators. One, it utilizes a lyophilized core antibody panel that allows clear identification of major compartments of the immune system and provides higher resolution into T Cell, B Cell, and other subsets. Lyophilization streamlines sample processing and eliminates the variability inherent in pipetting small volumes from a large numbers of individual antibody vials. Two, a two-tiered barcoding scheme assures that all donors and treatments are acquired together and that samples are organized into batches. This reduces the technical variation associated with the instrument and its operation. Three, a fully automated cloud-based analytics platform (Astrolabe) runs the same quality control, data cleaning, cell subset labeling, and unsupervised clustering over the entire data set. Taken together, the workflow provides a flexible framework that can be easily adapted to clinical trial immune monitoring or other large-scale experiments and greatly improve the quality, reproducibility, robustness and utility of mass cytometry data.

We leveraged this standardized workflow as part of a comprehensive screen to establish the expression of 350 surface markers across all major circulating immune subsets at single cell resolution. Acquisition of the entire expression dataset across three donors required more than a month of Helios operation and culminated in over 60 million events; one of the largest single mass cytometry datasets recorded to date.

Several quality control approaches were included in order to ensure the accuracy and quality of the antibody staining dataset. First, we employed a two-tier barcoding approach to minimize technical variability in performing the screen. The barcoded samples were deconvolved using an automated debarcoding approach that was directly compared and shown to perform comparably to manual debarcoding. Second, we used average overlap frequency (AOF) as a metric to evaluate the consistency of individual marker staining quality across all samples, which allows us to identify and address acquisition batch effects. Third, we used an automated approach to identify and label cell subsets, the accuracy of which was validated against manual gating of each of the analogous subsets, demonstrating high overlap and consistency between these approaches. Fifth, we performed the screen using three independent donor blood samples to allow for an evaluation of the biological reproducibility of individual marker expression profiles, and each donor presented a consistent and distinct cell subset profile across the entire experiment with both the frequencies of the major immune compartments and the intensities of their canonical markers showing low variability across the entire acquisition period. Finally, the reproducibility of the antibody expression profiles in our primary screen were further validated using a second independent screen performed using an additional donor. Taken together, these steps highlight the fidelity of the antibody staining resource. However, it is still important to note the limitations of this data set as a high-throughput screen; any findings require independent follow-up to confirm whether the reported expression patterns truly reflect hitherto unknown phenotypic diversity or may reflect specific biological or technical aspects of this screen. As an illustration of this approach, we used the screen to identify potential markers to characterize CD161+ MAIT cells, and then performed an independent experiment where we incorporated these markers as part of a single CyTOF panel. This allowed us to both independently validated the markers identified the screen and to further explore their co-expression patterns, confirming that CD161hi MAIT cells can be further characterized as being CD26hi, CD192hi, CD183low and CD57low.

In addition to screening marker expression patterns on fresh cells, we also introduced formaldehyde fixation as a treatment, thoroughly examining the influence that this standard perturbation could have on surface marker staining. When examining the effect of fixation on marker expression patterns, 173 out of 255 expressed markers had no change in their intensity. 65 gained some signal from fresh to fixed. We hypothesize that this gain is an artifact of the fixation protocol rather than a novel biological signal since it was subset agnostic and only affected some of the cells in each profiling subset. 17 markers lost their existing signal after fixation. In almost all cases, the loss of signal affected all expressing subsets. The one exception was CD22, where one expressing subset (basophils) lost the signal, while another (B Cells) did not. It has previously been suggested that the CD22 epitope on basophils is conformationally distinct from that on B cells (Toba et al., 2002). Our data provide further evidence suggesting a difference in the fixation sensitivity of the CD22 epitopes expressed on these two cell types.

The overall antibody staining data set is a powerful asset for immunologists seeking to investigate the immune system through the lens of less-explored markers and develop antibody panels to focus on specific cell subsets. To maximize the utility of this versatile qualitative resource, these results are fully accessible through an interactive website at https://www.astrolabediagnostics.com/antibody_staining_data_set. We included two aggregate statistics for each (marker, subset) combination: median anti-PE intensity and percent positive cells (which was calculated based on the background intensity available in the Blank LEGENDScreen well). In addition to interacting with the dataset through heat maps, survey aggregate statistics for their marker(s) and cell subset(s) of choice, the website allows investigators to delve deeper into the single-cell resolution and the relevant distributions. Overall, this dataset represents an accessible and unbiased resource for assessing potential expression of various markers over a large range of immune subsets in healthy individuals and surveying the statistics in the entire data set reveals intriguing signals for potential expression of less-studied markers. This study offers a valuable new resource to aid in the design of high dimensional antibody panels for immune monitoring studies, and further offers a template for a robust experimental workflow incorporating several components to ensure the accuracy and robustness of data generated using mass cytometry technology.

## Supporting information

Supplementary Table 1: Core Panel Antibodies

Supplementary Table 2: LEGENDScreen Antibodies

Supplementary Table 3: Astrolabe Labeling Hierarchy

Supplementary Table 4: Comparison of ASDS versus Validation

Supplementary Table 5: Antibody Panel for CD161 Validation

## 6 Author Contributions

EDA contributed to experiment design, analysis, and writing the manuscript. BL, PB and XVG contributed to sample acquisition and analysis. MM contributed to experiment design and writing the manuscript. AHR contributed to experiment design, sample acquisition and analysis, and writing the manuscript.

## 8 Acknowledgments

The authors would like to thank Shermineh Bradford, Geoffrey Kelly, Bhaskar Upadhyaya and the Human Immune Monitoring Center for technical assistance.

## 10. Data Availability Statement

All datasets acquired in the course of this study will be available on ImmPort (https://www.immport.org) upon manuscript publication.

## Footnotes

1 https://www.nih.gov/news-events/news-releases/nih-partners-11-leading-biopharmaceutical-companies-accelerate-development-new-cancer-immunotherapy-strategies-more-patients

