## Supplementary Material for "Development of a Comprehensive Antibody Staining Database using a Standardized Analytics Pipeline"

### 1 Supplementary Figures

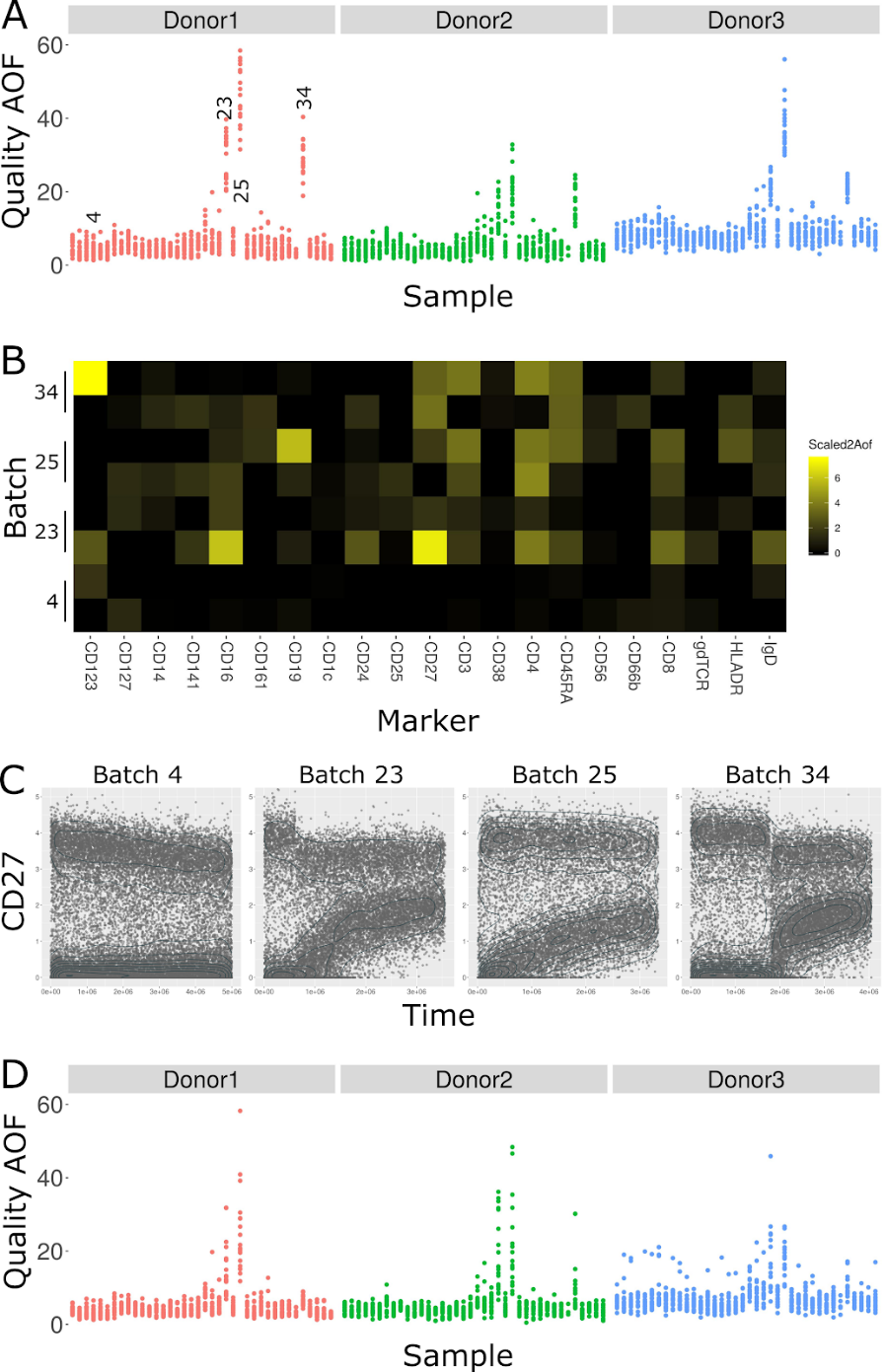

**Supplementary Figure 1. The average overlap frequency (AOF) identifies staining issues in three batches.** A: Scatter plot of the Quality^2^AOF in each sample. X-axis is batch, Y-axis is Quality^2^AOF, dots are samples color-coded by donors. Three of the batches (23, 25, and 34) showed higher Quality^2^AOF than others across all three donors in the batch. B: Heat map of Scaled^2^AOF in two samples out of each of the batches highlighted in A. While Scaled^2^AOF values are low for almost all markers in batch 4, there are several problematic markers in the other batches, notably CD123, CD16, CD19, and CD27. C: Scatter plot of CD27 staining in batches 4 (which passes AOF quality control), 23, 25, and 34. X-axis is acquisition time, Y-axis is CD27 intensity, dots are events. The contour plot (in jade) is event density. For batch 4, there is some CD27 signal degradation, but it does not interfere with downstream analysis. Batches 23, 25, and 34 each have significant increases in the negative modality which complicate identification of CD27+ versus CD27- cells. D: Scatter plot of Quality^2^AOF after gating on time in batches 23, 25, and 34. These batches were salvaged by filtering down to events acquired before the signal degradation.

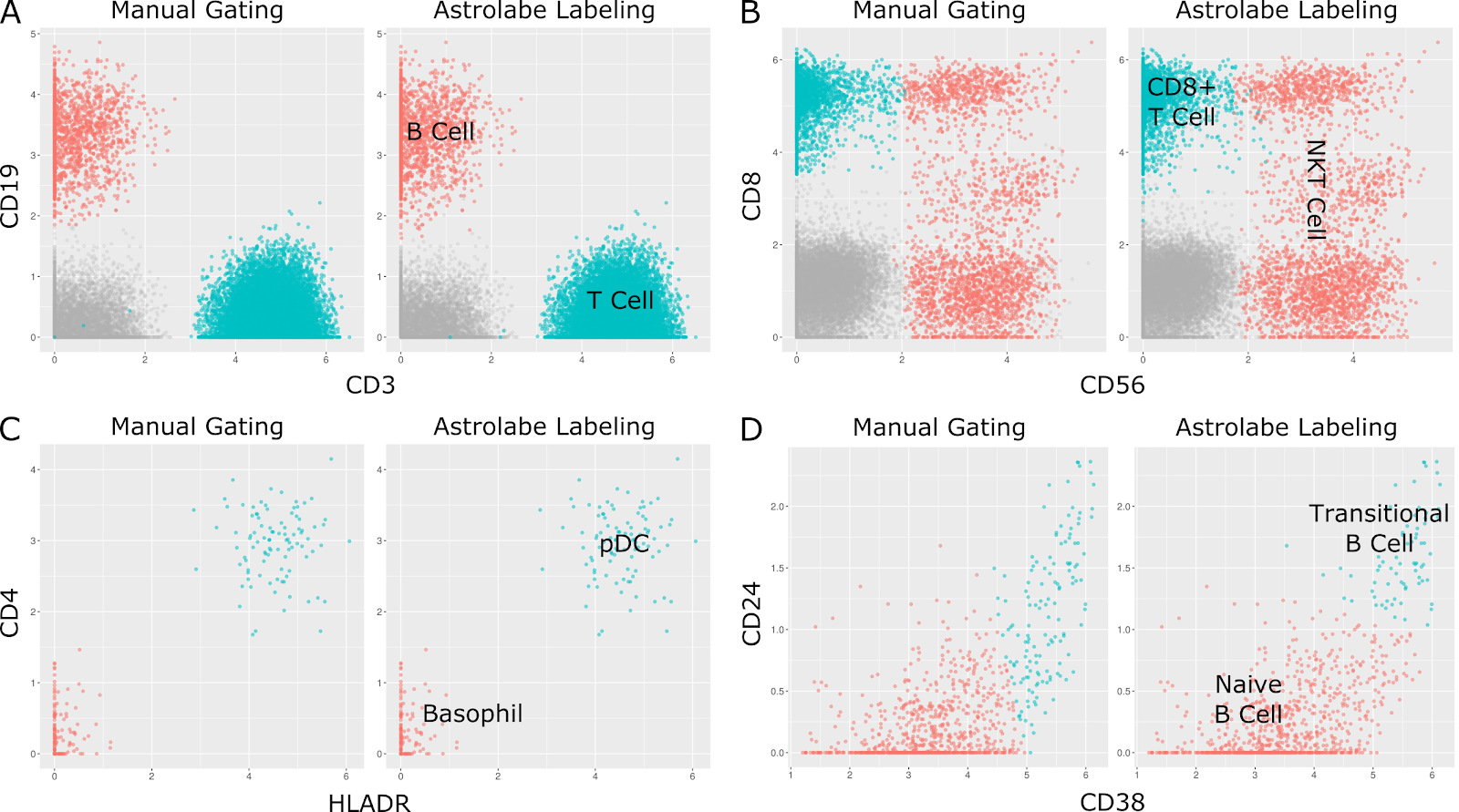

**Supplementary Figure 2. Comparison of biaxial plots between manual gating and Astrolabe labeling.** In each of these panels, dots are single cells. Left plot is manual gating, right plot is Astrolabe labeling. Color-coding is consistent between the two methods, and is indicated in the right plot. Grey dots do not belong to either of the subsets indicated. A: Separation of T Cells and B Cells based on CD3 and CD19. B: Within the T Cell compartment, separation of NKT Cells and CD8+ T Cells. C: Within the CD123+ compartment, separation of Plasmacytoid Dendritic Cells (pDCs) and Basophils. D: Within the CD27+ B Cell compartment, separation of Transitional B Cells and Naive B Cells. The two methods disagree on where to draw the CD38/CD27 threshold for Transitional B Cells. The manual gating operator chose a diagonal that includes CD24- cells while Astrolabe labels Transitional B Cells as CD38+ CD24+.

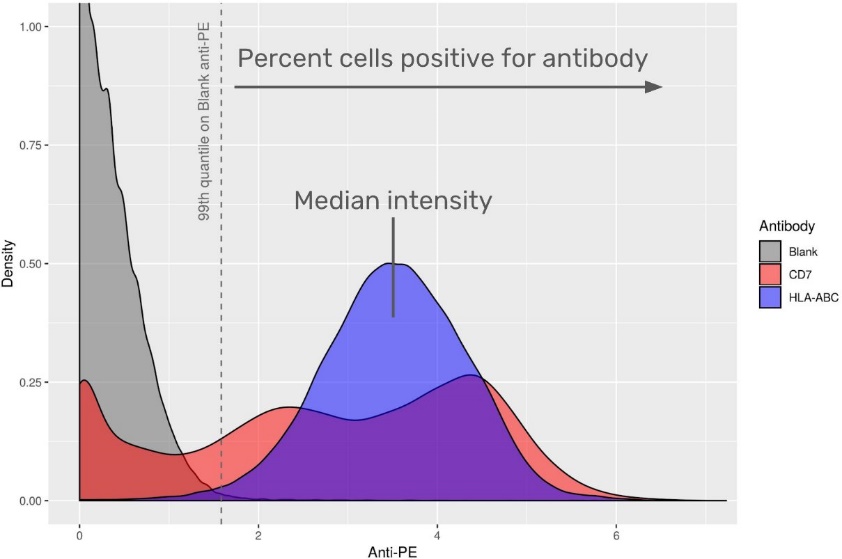

**Supplementary Figure 3. Calculation of aggregate statistics for the antibody staining data set.** Anti-PE intensity distribution for three antibodies: Blank (well A1 in plate 1 in the LEGENDScreen kit, which does not include a PE-conjugated antibody), CD7, and HLA-ABC. X-axis is anti-PE, Y-axis is density. Distributions are color-coded by antibody. The blank well provides the background intensity for staining. The heat map in Figure 4 includes two aggregate statistics for each (antibody, profiling subset) combination: percent positive and median intensity. Percent positive is the percent of subset cells which have anti-PE above above the 99th percentile on the blank well. Median intensity is the median anti-PE over all subset cells, regardless of whether their anti-PE is above or below the Blank 99th Percentile.

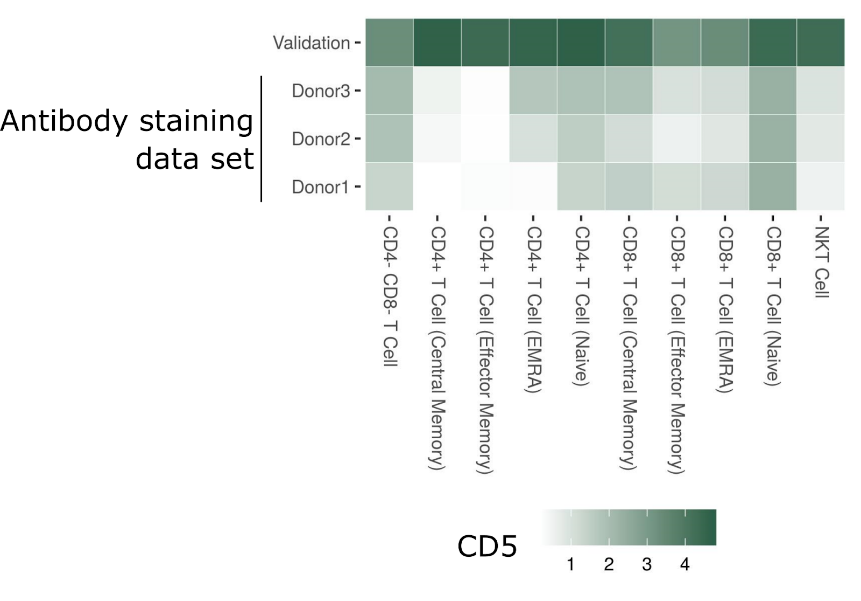

**Supplementary Figure 4. Comparing CD5 anti-PE intensity between the antibody staining data set and validation LEGENDScreen experiment.** Heat map of median CD5 intensity across different T Cell subsets. Columns are subsets, rows are samples (Donor1-3 are from the antibody staining data set). Tile intensity is median CD5 anti-PE intensity. The validation sample has strong CD5 expression across all T Cell subsets. However, for the donors in the antibody staining data set, expression is weaker, and is entirely missing from memory CD4+ T Cells and NKT Cells.

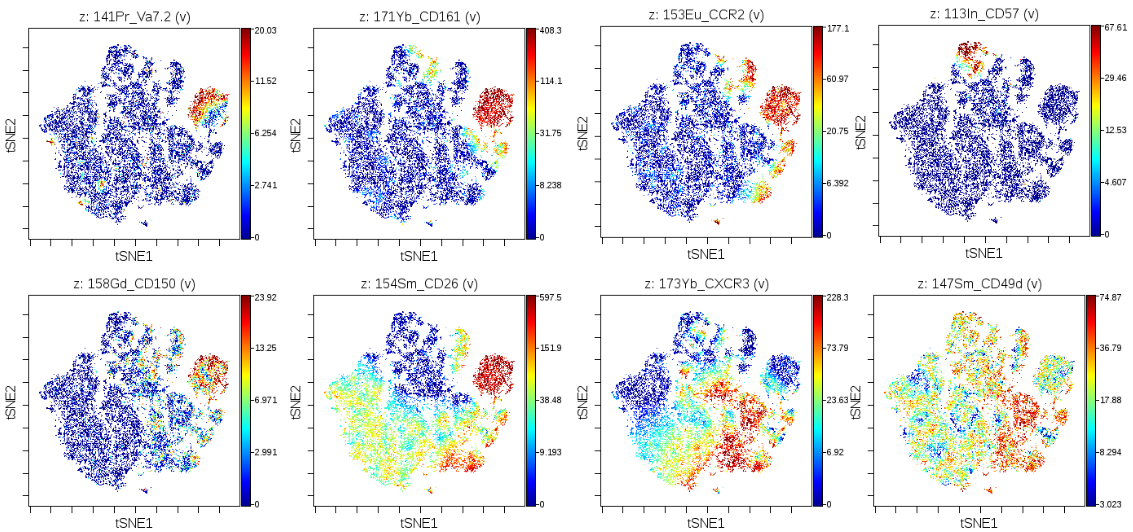

**Supplementary Figure 5. t-SNE maps of CD8+ T Cells, color coded by expression values of several in-panel markers.** The CD161+ island is located to the top right. The panels highlight the correct identification of the CD161+ CD8+ T Cells, and validate the expression of the markers flagged by the LEGENDScreen.

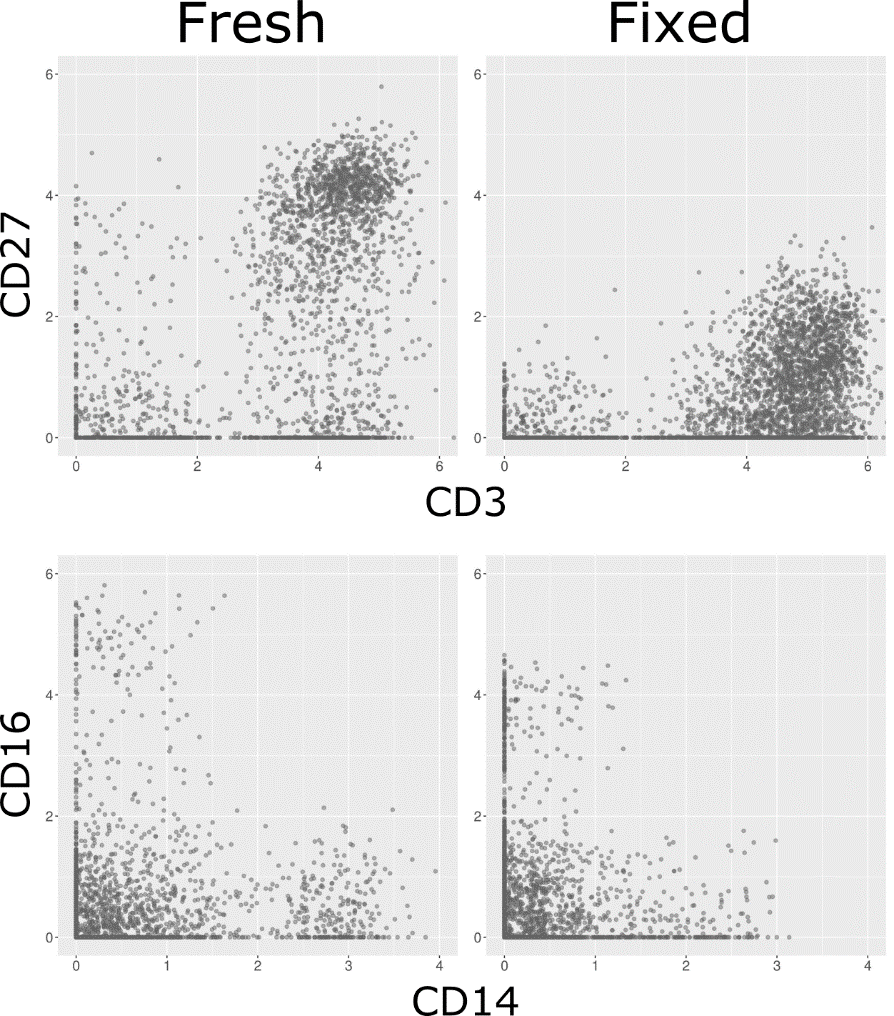

**Supplementary Figure 6. Loss of marker intensity in fixed samples.** Biaxial plots of four markers, before (left) and after (right) fixation. Each dot is a cell. Top: Loss of CD27 signal makes distinguishing CD27- and CD27+ T Cells challenging. Bottom: Loss of both CD14 and CD16 signal. While CD16+/- can still be distinguished, it is more challenging for CD14.

**Supplementary Table 1. List of antibodies, masses, and clones in the broad lyophilized panel.**

**Supplementary Table 2. List of antibodies and clones in the LEGENDScreen kit.**

**Supplementary Table 3. Astrolabe gating hierarchy.** The rules that were provided to the Ek’balam algorithm when labeling the cell subsets in the antibody staining data set. See “The Ek’balam algorithm” below for more details.

**Supplementary Table 4. Comparison of antibody staining data set and validation experiment.** Each row corresponds to a (Cell Subset, Antibody) combination. PerPos corresponds to percent positive cells in each donor and validation experiment. MedianAntiPe is median of anti-PE. Residual is calculated based on a linear regression between all data set and validation points.

**Supplementary Table 5. List of antibodies, masses, and clones in the CD161 validation.**

### 2 The Ek’balam algorithm

The Ek’balam algorithm receives as input a flow or mass cytometry sample and outputs a subset label for each cell in the sample. Labeling is done according to a table called a gating hierarchy, a list of rules that is manually curated by the user and maps marker combinations into subsets. The hierarchy used for the antibody staining data set is provided in Supplementary Table 1.

The hierarchy includes two types of subsets, Parent and Terminal subsets. Parent subsets can be further classified into additional subsets. For example, the “T Cell” subset can be split into “CD4+ T Cell”, “CD8+ T Cell”, etc. In the context of the Parent, these are called Child subsets. One special Parent subset is “Root”, which is the initial subset for all cells. Terminal subsets, such as “Basophil” or “gd T Cell” cannot be divided further.

The outline of Ek’balam is as follows:

- Label all cells as “Root”
- For each Parent subset that has cells labeled with it,
  - Define ParentCells as all cells labeled with Parent
  - Define ParentMarkers as the set of markers used by any of the Child rules under Parent
  - Repeat the following N times
    - Clustering: Cluster ParentCells over ParentMarkers
    - Marker Classification: Find all (Cluster, ParentMarker) combinations where Cluster is positive for ParentMarker
    - Matching: Match each Cluster to the Child rules under Parent. Cells in each Cluster get a vote for that Cluster’s Child
  - Vote: For each Child subset, find all ParentCells that have at least K votes for that Child and label them with that Child
  - Any ParentCell that does not have a Child subset is labeled as Parent_unassigned
- Stop when all cells belong to a Terminal subset

The following subsections describe the Clustering, Marker Classification, Matching, and Voting steps. Each section also details the parameters for that step and the default values used by Astrolabe in the antibody staining data set.

#### 2.1 Clustering

Generally speaking, any suitable clustering method could be used in the context of Ek’Balam. Astrolabe uses FlowSOM (citation needed) due to its high speed and accurate clustering (citation needed). In each iteration, ParentCells are clustered over ParentMarkers. Astrolabe uses a square SOM (where xdim = ydim). The value of SomDim is defined using the heuristic,

min{ceiling[sqrt(2^D^)], ceiling[sqrt(4 x N)]}

Where D is the number of ParentMarkers and N is the number of Child rules.

Briefly, the heuristic incorporates two considerations. One, we assume that each marker is positive or negative, leading to a total of 2^D^ combinations. Since we are using a square SOM, by setting xdim and ydim to ceiling[sqrt(2^D^)] we will get a number of clusters as close to 2^D^ as possible. Two, we ideally want one cluster for each Child. Since clustering algorithms often split subsets, so we relax that assumption to 4 x N (and again, ceiling[sqrt(4 x N)] due to the square SOM). However, we only need to fulfill one of these considerations, hence taking the minimum over both values.

#### 2.2 Marker Classification

The next part of the algorithm is classifying the Cluster marker profiles: which ParentMarker is positive for each Cluster. For a given ParentMarker, the Clusters which are positive for that marker are found using:

argmax _Partition ∈ Partitions , 0 < Threshold < max(ParentMarker)_ MCC(Partition, Threshold)

Where Partitions are all possible Cluster combinations and MCC is the Matthews Correlation Coefficient. In other words, we are looking for the combination of Clusters and the marker Threshold which maximizes the MCC. Instead of looking for an optimal solution, we use the following greedy algorithm for each marker:

- Order Clusters in increasing median marker intensity
- Start with an empty Partition
- For each Cluster in the list of Clusters,
  - Add the Cluster to Partition
  - Calculate the MCC using Partition and Thresholds of the 0%, 1%, 5%, 10%, and 20% percentiles over Cluster marker intensities
- Choose the Partition, Threshold combination with the higher MCC

Instead of testing all possible Partitions, we start with an empty Partition and add the Clusters one-by-one, according to their median marker intensity order. Additionally, instead of testing all possible Thresholds, in each iteration we test five possible Thresholds which are based on the latest Cluster to be added.

The above algorithm provides us with the list of Clusters which are positive for each ParentMarkers. By transposing this list we receive the list of ParentMarkers which are positive for each Cluster. This marker profile is the input for the next step, matching.

#### 2.3 Matching

In the matching step, each Cluster is matched to one of the Parent’s Child subsets using the marker profile for that Cluster and based on the Child rules. To demonstrate, here are the Child subsets that diverge from “Root” in the antibody staining data set gating hierarchy and their respective rules:

| **Parent** | **CellSubset** | **CD3** | **CD19** | **CD56** | **CD66b** | **CD14** | **HLADR** |
| --- | --- | --- | --- | --- | --- | --- | --- |
| Root | CM- | FALSE | FALSE | FALSE | FALSE | FALSE | NA |
| Root | T Cell | TRUE | FALSE | NA | NA | FALSE | NA |
| Root | B Cell | FALSE | TRUE | FALSE | NA | FALSE | NA |
| Root | CD14+ Monocyte | FALSE | FALSE | FALSE | NA | TRUE | NA |
| Root | NK Cell | FALSE | FALSE | TRUE | FALSE | FALSE | FALSE |
| Root | Granulocyte | FALSE | FALSE | FALSE | TRUE | FALSE | NA |

In this example, six markers are considered when labeling “Root” cells: CD3, CD19, CD56, CD66b, CD14, and HLADR. Any cluster that is CD3+, CD19-, and CD14- will be matched to “T Cell”. The values of CD56, CD66b, and HLADR do not influence that matching. Likewise, any cluster that is CD3-, CD19-, CD56+, CD66b-, CD14-, and HLADR- will be matched to “NK Cell”, and so on.

There are two special cases for matching:

- Any clusters that don’t match any of the rules will be matched to the default “Parent_unassigned” rule (in this case, “Root_unassigned”)
- Clusters that match two or more rules will lead to an error. The hierarchy should be designed in such a way to avoid matching to more than one rule

In the next iteration, cells labeled “CM-”, “T Cell”, “B Cell”, “CD14+ Monocyte”, and “NK Cell” will each be clustered separately and further labeled. Cells labeled “Granulocyte” and “Root_unassigned” will not be clustered, since this is a Terminal label.

Here are the subsets that diverge from “B Cell”:

| **Parent** | **CellSubset** | **ClusterOn** | **CD27** | **CD38** | **CD24** |
| --- | --- | --- | --- | --- | --- |
| B Cell | B Cell (CD27+) | CD19 | TRUE | NA | NA |
| B Cell | B Cell (Transitional) | CD19 | FALSE | TRUE | TRUE |
| B Cell | B Cell (CD24- CD27-) | CD19 | FALSE | NA | FALSE |

Four markers are considered for the “B Cell” subset: CD27, CD38, CD24, and the additional CD19 that appears under ClusterOn. The ClusterOn column can include any number of markers which will be added to ParentMarkers (and therefore to the Clustering step). After clustering over these four markers, CD27+ Clusters will be matched to “B Cell (CD27+)”, CD27- CD38+ CD24+ Clusters to “B Cell (Transitional)”, and CD27- CD24- Clusters to “B Cell (CD24- CD27-)”. All other “B Cell” cells will be labeled “B Cell_unassigned”.

#### 2.4 Voting

Many of the algorithms used for clustering in the context of cytometry data include a stochastic element, usually in the form of a random seed, which influences the algorithm output. It is very common to run an algorithm multiple times and get different cluster assignments. While methods exist for finding the consensus over multiple clustering runs (citation needed), they are agnostic of the context where the data was acquired. Ek’balam builds on top of the gating hierarchy to achieve a stronger consensus mechanism through voting.

For each Parent subset, the Clustering, Marker Classification, and Matching steps are run multiple (N) times. Each such iteration contributes one vote for each cell. Once they are all done, cells that have a number of votes over a given threshold (K) get labeled with a Child rule. All other cells are labeled as Parent_unassigned.

For the Astrolabe implementation, N is defined as 25 if SomDim is g.t.e 5, and 10 otherwise. K is defined as floor(N / 2).

### 3 Cell subset profiling

Profiling is an extension to the Ek’Balam algorithm portrayed above. Ek’Balam terminates once every cell is labeled with a Terminal subset. Profiling adds one or more levels of clustering, marker classification, and matching. The new levels are not supplied by the user. Instead, they are generated in an unsupervised fashion using data from all of the samples in the experiment.

The profiling algorithm operates on each cell subset separately. For a given cell subset, the outline is as follows:

- Define Markers as every marker which was not used in the hierarchy reaching to this cell subset
- For each sample,
  - Cluster subset using the same parameters as Ek’Balam
  - Calculate the maximum MCC for each Marker
- Calculate the median maximum MCC for each Marker over all samples and order Markers in decreasing median maximum MCC
- For each Marker in the list of Markers,
  - Calculate frequency of cells positive for this marker over all samples
  - If the frequency is greater than 2%, add this Marker to the profiling level of the hierarchy
  - Otherwise, stop

The algorithm is designed around balancing two competing interest. First, we would like to see as many interesting markers for a given subset, where we define interesting as consistent separation between clusters across all samples. However, we would like to avoid a situation where some markers are too small to be biologically relevant, defined as at least 2% of the total cells in the sample.

The result is an iterative algorithm which repeatedly splits the cell subset according to MCC order. However, as soon as we end up splitting the subset too small, the algorithm stops. The resulting level or levels are then appended to the Ek’Balam algorithm, as Child subsets of each (previously) Terminal subset.
